# The interferon-rich skin environment regulates Langerhans cell ADAM17 to promote photosensitivity in lupus

**DOI:** 10.1101/2021.08.18.456792

**Authors:** Thomas M. Li, Victoria Zyulina, Ethan S. Seltzer, Marija Dacic, Yurii Chinenov, Andrea R. Daamen, Keila R. Veiga, Noa Schwartz, David J. Oliver, Jose Lora, Ali Jabbari, Yong Liu, William D. Shipman, William G. Ambler, Sarah F. Taber, Karen B. Onel, Jonathan H. Zippin, Mehdi Rashighi, James G. Krueger, Niroshana Anandasabapathy, Inez Rogatsky, Carl P. Blobel, Peter E. Lipsky, Theresa T. Lu

## Abstract

**Background:** The autoimmune disease lupus erythematosus (lupus) is characterized by photosensitivity, where even ambient ultraviolet radiation (UVR) exposure inflames skin. Beneficial effects of anifrolumab (anti-interferon α/breceptor (anti-IFNAR)) on lupus skin disease support a pathogenic role for IFN-I, but mechanistic understanding is limited. We have shown that Langerhans cell (LC) dysfunction contributes to photosensitivity. Healthy LCs act via a disintegrin and metalloprotease 17 (ADAM17) to release epidermal growth factor receptor (EGFR) ligands that limit UVR-induced keratinocyte apoptosis and photosensitivity. However, LC ADAM17 activity is reduced in non-lesional lupus model skin, and data point to reduced LC-mediated protection in human lupus. Here, we asked about the role of the IFN-rich lupus skin environment in LC dysfunction and the implications of this regulation for photosensitivity.

**Methods:** Gene expression patterns in non-lesional skin from human lupus and multiple murine models were examined. We used MRL/lpr, B6.Sle1yaa, and imiquimod models of lupus in *in vivo* studies to assess the role of IFN-I in LC ADAM17 dysfunction and photosensitivity.

**Results:** We show a shared IFN-rich environment in non-lesional skin across human and murine model systems, that IFN-I inhibits LC ADAM17 activity, and that anti-IFNAR in lupus models restores LC ADAM17 function and reduces photosensitivity in EGFR and LC ADAM17-dependent manners. Reactive oxygen species (ROS) can mediate ADAM17 activity, and we show reduced LC ROS expression in lupus models that is restored by anti-IFNAR.

**Conclusions:** Our findings suggest that IFN-I promotes photosensitivity by causing LC ADAM17 dysfunction and that anifrolumab ameliorates lupus skin disease at least in part by restoring LC function. This work provides insight into IFN-I-mediated disease mechanisms, LC regulation, and a mechanism of action for anifrolumab in lupus.

## INTRODUCTION

The autoimmune disease lupus erythematosus (lupus) manifests in part by photosensitivity, a sensitivity to ultraviolet radiation (UVR) whereby even ambient sunlight can trigger the development of inflammatory skin lesions. Patients with lupus skin disease, or cutaneous lupus erythematosus (CLE), can also have the systemic form of the disease, systemic lupus erythematosus (SLE), characterized by inflammation and damage to additional organs such as the kidneys, brain, and heart. In SLE patients, photosensitive skin responses can be associated with worsening systemic autoimmunity and end organ damage. A prominent interferon (IFN) signature indicative of exposure to type I interferon (IFN-I) is found in circulating cells and tissues in lupus (Baechler et al., 2003; Bennett et al., 2003; Catalina et al., 2019; Der et al., 2019; Jabbari et al., 2014; Kirou et al., 2004), and the development and recent FDA approval of anifrolumab (anti-IFNAR1) for SLE highlights the importance of IFN-I in disease pathogenesis. Anifrolumab is especially efficacious for skin disease in SLE patients, consistent with reduced skin lesions in IFNAR-deficient SLE model mice (Nickerson et al., 2013) and an IFN-I signature in both lesional and non-lesional skin (Billi et al., 2022; Der et al., 2017; Der et al., 2019; Reefman et al., 2008; Stannard et al., 2017). These findings suggest a prominent role for IFN-I in lupus skin disease, but mechanisms of IFN-I-mediated pathogenesis remain poorly understood.

We recently reported that Langerhans cells (LCs) can limit UVR-induced skin inflammation by expressing a disintegrin and metalloprotease 17 (ADAM17) which releases membrane-bound epidermal growth factor receptor (EGFR) ligands that then preserve epidermal integrity via EGFR stimulation of keratinocytes (Shipman et al., 2018). Two distinct photosensitive SLE mouse models showed reduced LC *Adam17* mRNA which corresponded to reduced LC ADAM17 sheddase activity, and photosensitivity was reduced by topical EGFR ligand application. Together, these data suggested that the lack of LC ADAM17-dependent EGFR-ligand sheddase activity contributes to photosensitivity. Non-lesional skin of SLE patients showed reduced LC numbers along with reduced epidermal EGFR phosphorylation, further supporting the potential for LC defects to contribute to photosensitivity in lupus. The factors that lead to LC dysfunction, however, remain to be fully elucidated.

Here, we test the hypothesis that the high IFN-I environment present in even non-lesional lupus skin promotes LC ADAM17 dysfunction to contribute to photosensitivity. We build upon human data showing an IFN-I signature in non-lesional skin of CLE patients and establish that the photosensitive MRL/lpr and B6.Sle1yaa models share IFN-I-related and other gene expression modules with human skin. The inducible photosensitive imiquimod (IMQ) model mice also upregulated IFN-associated genes in non-lesional skin. IFN-I is sufficient to downregulate healthy murine and human LC ADAM17 activity, and anti-IFN-I receptor (anti-IFNAR) treatment of lupus mouse models restores LC ADAM17 activity and reduces photosensitivity in an EGFR and LC ADAM17-dependent manner. Lastly, reactive oxygen species (ROS) are known to promote ADAM17 activity (Singh et al., 2009), and we show that UVR-induced LC ROS are reduced in lupus model mice while anti-IFNAR restores ROS expression. Together, our results establish that multiple photosensitive murine lupus models are similar to human lupus patients in demonstrating IFN-I signatures in non-lesional skin and put forth a mechanism whereby IFN-I contributes to photosensitivity at least in part by inhibiting LC ADAM17 activity. These data provide insight into IFN-I-mediated contributions, delineate a driver of LC dysfunction, and suggest that restoration of LC ADAM17 function is a mechanism of action for anifrolumab in lupus skin disease.

## RESULTS

### Non-lesional skin in lupus model mice share an IFN-I signature and other functional gene expression modules with human lupus skin

As we are interested in the mechanisms that predispose to photosensitivity and to the LC defects seen in lupus mouse models and human SLE skin (Shipman et al., 2018), we sought to better understand the environment in such non-lesional skin. Recent studies have suggested a high IFN-I environment in non-lesional skin of lupus patients based on single cell RNAseq of keratinocytes from SLE patients and skin from CLE patients (Billi et al., 2022; Der et al., 2017; Der et al., 2019), CLE keratinocyte IFN-κ expression (Stannard et al., 2017), and upregulation of IFN-stimulated genes (ISGs) such as *MX1* on tissue sections in SLE and incomplete SLE patients (Lambers et al., 2019; Reefman et al., 2008). Consistent with these findings, our recent analysis of publicly available bulk transcriptomic datasets from non-lesional DLE skin using Gene Set Variation Analysis (GSVA), an unsupervised gene set analysis approach, showed a proportion of samples with upregulated IFN gene set expression when compared to healthy controls, although this effect was modest when compared to the upregulation in lesional skin (Martínez et al., 2022). Here, we examined an additional non-lesional skin dataset from a CLE cohort with the discoid form of CLE (DLE) and also non-lesional skin from multiple lupus models to understand shared pathways.

The non-lesional skin has not been previously analyzed and comes from a DLE cohort whose lesional skin had been shown to have a Th1 signature on microarray when compared to healthy controls and psoriatic skin (Jabbari et al., 2014). This was among the lesional skin cohorts that was shown by GSVA to express the IFN gene set at high levels (Martínez et al., 2022). Normal (ie healthy subject) and non-lesional skin exhibited significantly different expression profiles (Figure 1A and Table S1). Differential pathway analysis via Quantitative Set Analysis for Gene Expression (QuSAGE) showed that IFN-I and IFN-γ pathways were among the most upregulated in the non-lesional skin compared to normal controls (Figure 1B and Table S2). The IFN-I pathway was comprised of several upregulated IFN-stimulated transcription factors, including IRF 1, 2, 7, 8, and 9 along with multiple IRF targets (ISG15, IFIT1, IFIT3, MX1, MX2) (Figure 1C-D, S1A). Notably, *CD207*, the gene encoding Langerin and a lineage marker of LCs, was downregulated (Figure 1C), consistent with accumulating data on LC dysfunction in lupus (Billi et al., 2022; Martínez et al., 2022; Shipman et al., 2018). A supervised approach to analyzing normal control versus non-lesional DLE skin gene expression data, then, yielded results consistent with other cohorts showing upregulation of IFN pathways and LC dysfunction.

**Figure 1.**
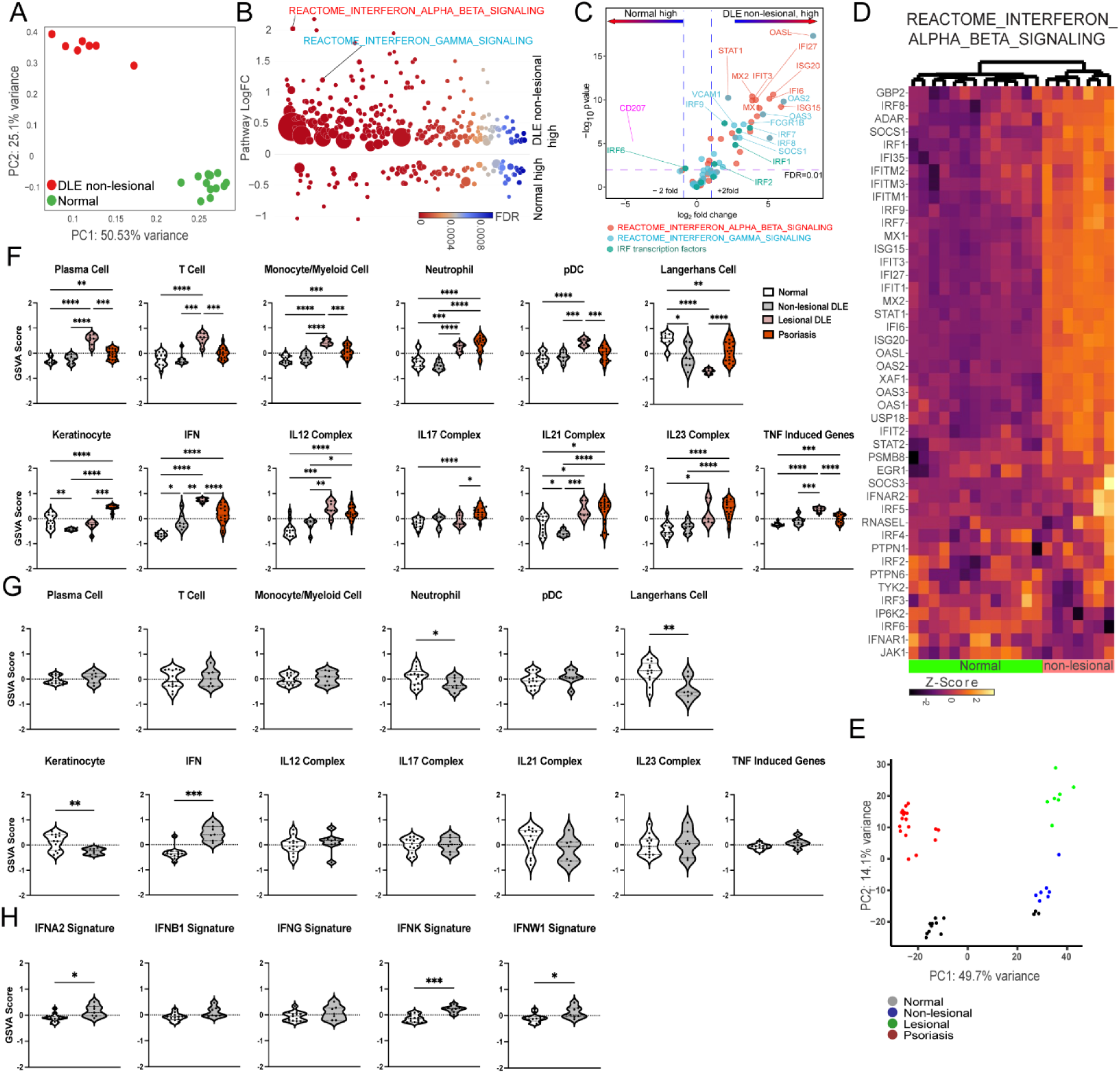
Analysis of a new cohort reinforces a model of an IFN-rich environment and LC and keratinocyte dysfunction in non-lesional DLE skin. (A-H) Microarray analysis of gene expression from non-lesional skin of DLE patients (n=7) and healthy controls (n=13). In (E-F), samples were analyzed with lesional DLE and psoriasis (PSO) skin samples that were collected as part of the same study and previously published (Jabbari et al., 2014). (A,E) Principal component analysis (PCA) using top 250 genes. (B) Differentially expressed pathways were determined using QuSAGE pathway analysis against Molecular Signatures Database (MSigDB). (C) Volcano plot of differentially expressed genes. Genes from IFN α/β (red), IFNγ (blue) and IRF transcription factor (green) pathways are marked. CD207 (pink) is additionally marked. (D) Heatmap of gene expression in the IFN α/β signaling pathway. (F, G) Gene Set Variation Analysis (GSVA) of gene sets relevant to lupus (Martínez et al., 2022). (H) GSVA using gene sets comprising specific IFN sub-types. (F-H) *p<0.05, **p<0.01, ***p<0.001, ****p<0.0001 by unpaired t-test.

We next performed GSVA using previously defined lupus-relevant gene sets (Martínez et al., 2022) (Table S3). Analysis of the non-lesional samples along with the previously analyzed lesional and psoriatic samples (Jabbari et al., 2014) reiterated the GSVA results of multiple datasets. The gene expression profiles of the four conditions were significantly different from each other (Figure 1E). Lesional DLE was characterized by enrichment of immune cell gene sets including plasma cells, T cells, monocyte/myeloid cells, neutrophils, and plasmacytoid dendritic cells (pDCs) in conjunction with de-enrichment of LCs when compared to normal controls (Figure 1F). Lesional DLE samples were also enriched for inflammatory cytokine gene sets including IL12, IL21, IL23, and TNF as well as a highly upregulated IFN gene signature as previously seen in lesional DLE skin (Figure 1F) (Martínez et al., 2022). Non-lesional skin also showed upregulation IFN when compared to healthy controls, although this change was dwarfed when compared to enrichment in lesional skin (Figure 1F). GSVA analysis of just healthy controls and non-lesional skin in the absence of the high IFN enrichment from lesional skin revealed clear upregulation of the IFN gene set in non-lesional skin as well as de-enrichment of LC and keratinocyte gene sets (Figure 1G). These results were consistent with our previous GSVA analyses of other non-lesional cohorts (Martínez et al., 2022), suggesting along with direct cellular analyses (Billi et al., 2022; Sarkar et al., 2018; Shipman et al., 2018) a loss of epidermal (LC and keratinocyte) function in the context of an IFN-rich proinflammatory environment. The IFN gene set upregulation in non-lesional skin was driven by IFNα, IFNκ, and IFNω gene set upregulation (Figure 1H). GSVA of these previously unanalyzed non-lesional DLE skin samples, then, reinforced previous gene expression analyses and revealed an IFN-I signature in this cohort, all pointing to perturbed epidermal and skin homeostasis even in non-lesional skin.

We examined non-lesional ear skin from MRL/lpr and B6.Sle1yaa SLE mouse models by RNAseq. The photosensitive MRL/lpr model has a *Fas* gene mutation and develops a lupus-like phenotype especially well on the MRL and but not the B6 genetic background. It has reduced LC ADAM17 sheddase function and mRNA, and topical EGFR ligand supplementation ameliorated photosensitivity (Shipman et al., 2018). Although the differences in gene expression between MRL/lpr mice and control MRL-MpJ (MRL/+) mice by RNAseq were less dramatic than those in the human cohort (Figures 1A-D and 2A-D) (Table S4), QuSAGE showed that the IFN-α/β and IFN-γ pathways were among the most highly expressed in MRL/lpr mice (Figure 2B and Table S5). Consistent with the activation of IFN pathways, several IRF transcription factors and their targets were expressed at higher levels in MRL/lpr mice (Figure 2C-D and Figure S1B).

**Figure 2.**
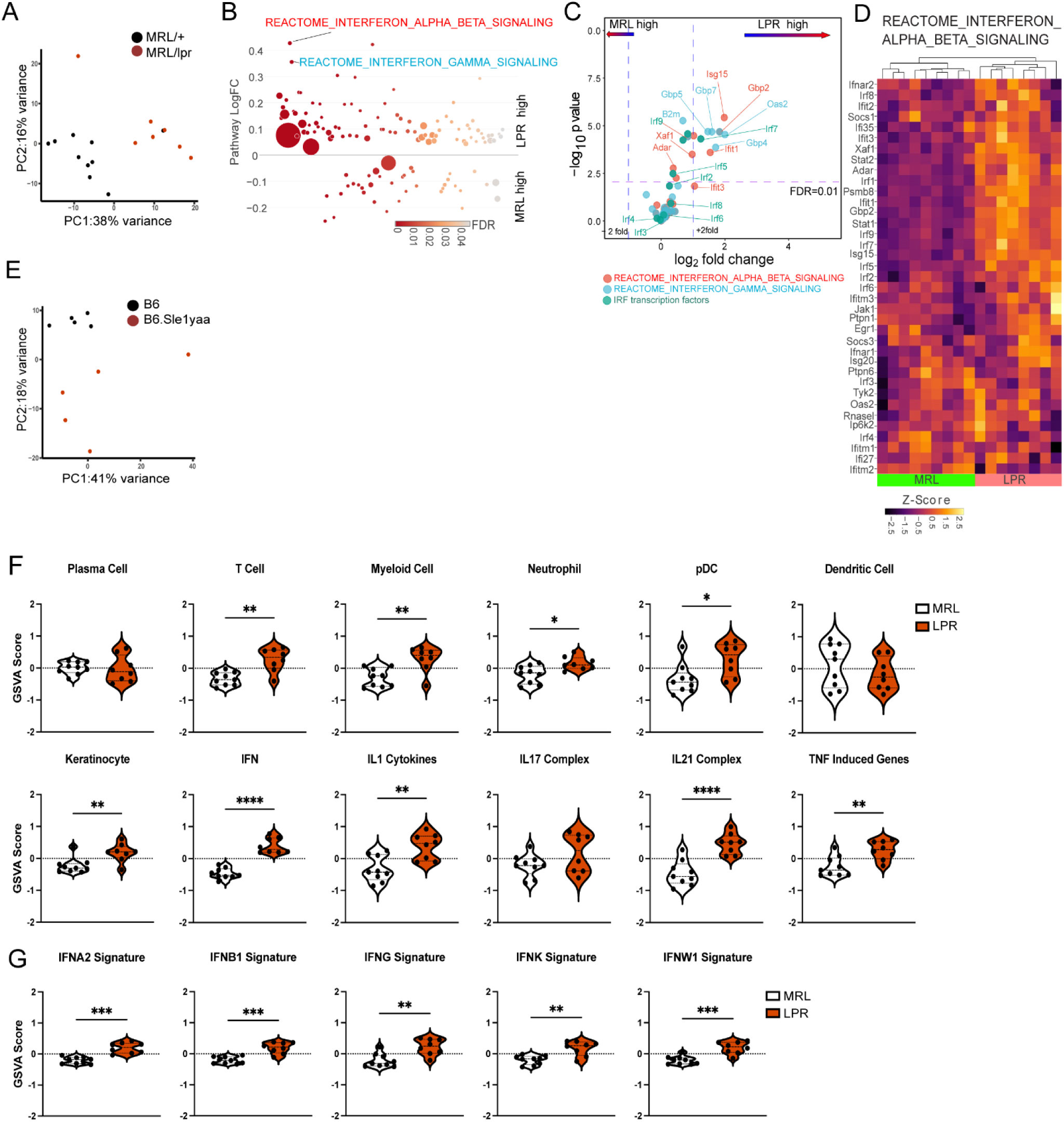
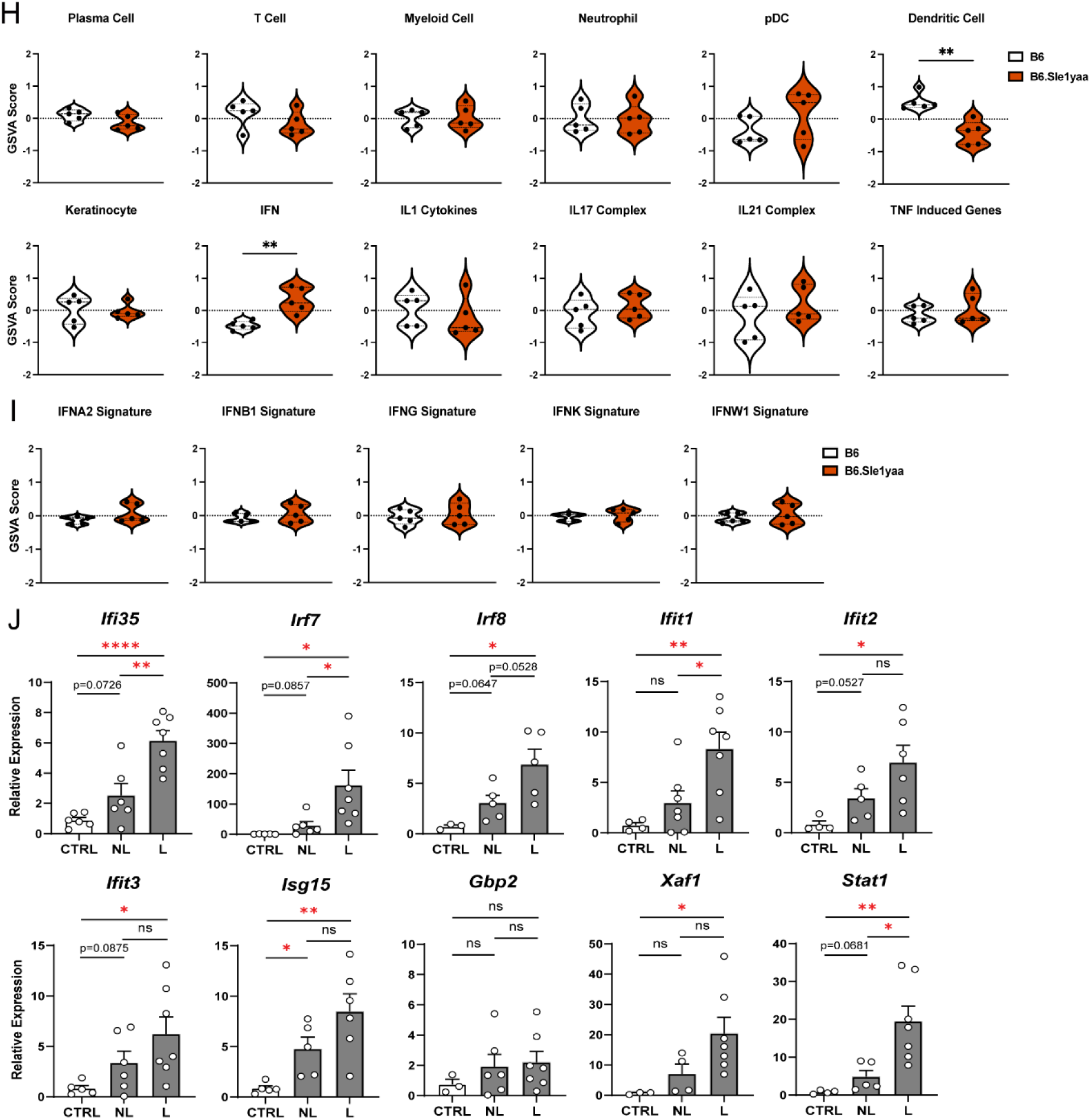
Non-lesional skin of multiple photosensitive murine SLE models recapitulate gene expression patterns of human DLE non-lesional skin. (A-I) Non-lesional skin gene expression from MRL/lpr (LPR) and control MRL/+ (MRL) mice (A-D, F-G) and B6.Sle1yaa and control B6 mice (E, H-I) were analyzed by RNA sequencing. (A, E) PCA using top 250 genes. (B) Differentially expressed pathways determined by QuSAGE pathway analysis against MSigDB. (C) Volcano plot of differentially expressed genes. Genes from IFN α/β (red), IFNγ (blue) and IRF transcription factor (green) pathways are marked. (D) Heatmap showing gene expression in the IFN α/b signaling pathway. (F, H) GSVA of gene sets relevant to lupus, adapted for murine models (Kingsmore et al., 2021). (G, I) GSVA of gene signatures specific to distinct IFN subtypes. (J) Imiquimod (IMQ) model non-lesional and lesional ears and control mice ears were analyzed by qPCR of select IFN-related genes that are upregulated in non-lesional skin of human DLE and MRL/lpr and B6.Sle1yaa models. (A, E-J) Each symbol represents one mouse (A, E-I) or one ear, with non-lesional and lesional ears coming from the same mice (J). (F-J) *p<0.05, **p<0.01, ***p<0.001, ****p<0.0001 by unpaired t-test.

The B6.Sle1yaa model is driven by the Sle1 lupus susceptibility locus derived from lupus-prone NZB2410 mice in combination with the Y chromosome autoimmune accelerator locus whose effects are attributed to *Tlr7* duplication (Pisitkun et al., 2006; Subramanian et al., 2006). These mice are photosensitive and have reduced LC *Adam17* mRNA expression (Shipman et al., 2018). As expected, compared to B6 controls, B6.Sle1yaa mice expressed elevated levels of several lupus related genes from the Yaa locus including Tlr7 (Figure S1C). Although differences between the B6 and B6.Sle1yaa transcriptomes were modest (Figure 2E)(Table S6), several genes involved in the regulation of and response to IFN were expressed at higher levels, including *Irf7, Irf9, Isg15, Xaf1, Selp,* and *Ube2l6* (Figure S1D). Although LC numbers were not different at 10 months of age when this model is fully diseased (Shipman et al., 2018), *Cd207* expression in younger (5 month old) diseased mice was downregulated when compared to B6 controls (Figure S1D), similar to the findings in CLE skin (Figure 1C). By supervised gene set analysis, both the MLR/lpr and B6.Sle1yaa murine models showed similarities to human non-lesional skin.

By GSVA of lupus-relevant murine gene sets (Table S7), parallels between the MLR/lpr and B6.Sle1yaa murine models and human lupus were even more dramatic. MRL/lpr mice showed increased enrichment of a number of inflammatory cell and cytokine gene sets including T cells, myeloid cells, neutrophils, pDCs, IFN, IL1, IL21, and TNF (Figure 2F). However, in contrast to human DLE, non-lesional skin of MRL/lpr mice was enriched for the keratinocyte gene signature. LC numbers in the skin of MRL/lpr mice are not downregulated when compared to MRL/+ controls (Shipman et al., 2018), which was reflected in the absence of a decreased dendritic cell signature. The IFN gene set upregulation was broad, with significant enrichment of gene signatures induced by stimulation from multiple IFNs (Figure 2G). B6.Sle1yaa mice showed less dramatic differences than the MRL/lpr model, but also showed upregulation of the IFN gene set (although there was not a significant contribution from any one IFN signature) and downregulation of the DC gene set, potentially reflective of decreased LC function (Figure 2H-1). Together, our results showed a number of parallels to the human system, including an IFN signature in these lupus models.

We also tested for expression of ISGs in the imiquimod (IMQ) lupus model. This model is induced by 4-5 weeks of topical IMQ, a TLR 7/8 agonist, resulting in autoantibody production, enlarged spleen and lymph nodes, nephritis, and photosensitivity characteristic of SLE (Ambler et al., 2022; Goel et al., 2020; Yokogawa et al., 2014). The model can be induced in B6 mice, allowing for the use of transgenic models to assess for effects on lupus features without having to cross onto a different genetic background. We applied IMQ to the right ear to assess ISGs in both the painted right “lesional” and unpainted left “non-lesional” ears. We performed qPCR for a set of transcripts (*Gbp2, Ifi35, Ifit1, Ifit3, Ifit2, Irf7, Irf8, Isg15, Stat1,* and *Xaf1)* that were upregulated across human and murine non-lesional skin datasets using our datasets (Figures S1A, B, D) and that in (Billi et al., 2022) (Figure 2J). Most of the genes we examined showed upregulation in the “lesional” ear, including transcripts for IRF target genes *Irf7, Irf8* and *Stat1*, and the apoptosis marker *Xaf1* potentially reflecting tissue damage along with an IFN-rich environment. Several transcripts also showed trends toward upregulation in non-lesional skin when compared to mice that were not treated with IMQ (Figure 2J). Notably, *Isg15*, the only transcript that we examined that was upregulated across all the datasets, was also upregulated in the non-lesional ears of the IMQ mice. ISG15 has been shown to limit over-amplification of IFN-I effects, and its deficiency predisposes to skin inflammation among other findings (Martin-Fernandez et al., 2020; Zhang et al., 2015), suggesting that upregulation of *Isg15* may reflect an attempt to minimize IFN-I-mediated inflammation. These results showed that non-lesional skin from human lupus and 3 murine models were characterized by upregulation of IFN-associated transcripts, albeit at lower levels than in lesional skin.

Turning our attention to common pathways in human and murine lupus non-lesional skin, we performed ranked Gene Set Enrichment Analysis (GSEA) separately for DLE and MRL/lpr datasets using negative log-transformed p-values as a ranking variable, constructed pathway similarity networks, and compared selected pathways (FDR<0.005) in humans and mice. Consistent with QuSAGE analysis (Figure 1B and 2B), IFN-related pathways were enriched in human DLE and MRL/lpr non-lesional skin (nodes 3, 4, 8), along with several other pathways often associated with lupus (Figure 3A): T helper pathway (nodes 26); natural killer cell-mediated cytotoxicity (nodes 28); Toll-like receptor pathways (nodes 36,46); JAK-STAT signaling pathway (node 43); and multiple pathways associated with complement activation (nodes 6, 23, 29, 34, 52, 53, 58, 61, 62, 64, 75, 88, 94, 95). For the full list of pathways identified by ranked GSEA, see Table S8. Direct comparison of differentially expressed genes in DLE and MRL/lpr skin (Figure 3B) revealed a gene signature typical of nucleic acid-driven interferonopathies dominated by IFN-stimulated transcription factors (IRF1,3,9 and STAT1), IFN-induced viral restriction factors (BST2, SAMHD1, DDX60, IFIH1, ZBP1), neutrophilic markers (HMOX1, S100A8, S100A9, LCN2, RSAD2), and apoptosis-related genes (XAF1, TNFSF10). Several keratins (KRT16, 6B and 34) that are rapidly induced upon skin damage and, in turn, downregulate inflammatory responses (Lessard et al., 2013), were highly expressed in both human CLE and MRL/lpr skin, likely reflecting disturbed epidermal homeostasis.

**Figure 3.**
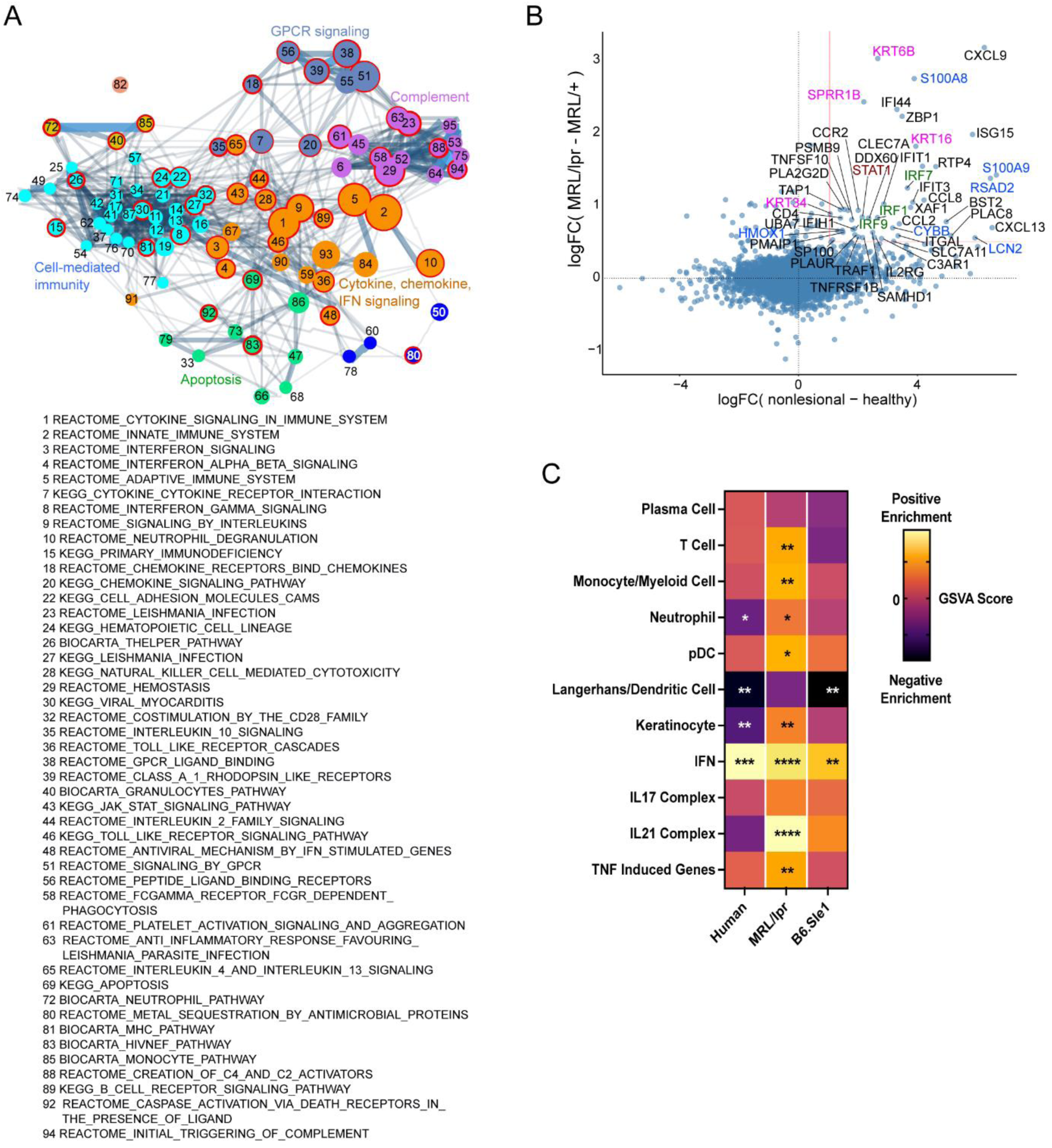
Analysis of human and murine lupus models show shared gene expression pathways. (A) Pathway similarity network built for the most enriched pathways (adjusted p<0.005) in DLE non-lesional skin determined by ranked GSEA (complete list in Table S8). Colors indicate network community clusters identified by the walktrap algorithm. Red borders indicate pathways that were also enriched in LPR mice (listed). (B) Comparison of gene expression changes in DLE vs MRL/lpr mice. (C) Heatmap of GSVA scores for shared gene sets between non-lesional skin from DLE patients, LPR mice, and B6.Sle1yaa mice. Asterisks indicate significant differences in GSVA scores compared to controls for each dataset.

Comparisons of the GSVA analyses of non-lesional skin among the datasets (Figure 3C) echoed the GSEA analysis in emphasizing the prominent enrichment of IFN in human CLE and both MRL/lpr and B6.Sle1yaa lupus-prone mice. In addition, de-enrichment of the LC/DC gene set was present in both human DLE and B6.Sle1yaa mouse non-lesional skin (Figure 3C). Multiple approaches to analyze gene expression of non-lesional skin from human DLE and murine lupus models, then, showed upregulation of IFN-related transcripts and disturbance of LC and epidermal homeostasis. These data are consistent with the idea that non-lesional skin is primed for photosensitive responses and that LCs sit within an IFN-rich environment in both human lupus and multiple murine models that may cause LC dysfunction.

### IFN-I inhibits LC ADAM17 function

We next examined the effects of IFN-I on LC ADAM17 function and expression. As previous (Shipman et al., 2018), we assessed LC ADAM17 function by quantifying UVR- induced cell surface TNFR receptor 1 (TNFR1) shedding. We confirmed that UVR treatment performed on a mixture of digested epidermal cells induced LC loss of surface TNFR1 in an LC ADAM17-dependent manner (Figure S2A-B), similar to our results with isolated LCs (Shipman et al., 2018). IFN-κ, an IFN-I that is highly expressed by lupus keratinocytes (Tsoi et al., 2019), reduced LC ADAM17 activity by nearly 40% (Figure 4A). Using a validated anti-murine ADAM17 antibody (Lora et al., 2021) (Figure S2A), we observed that IFN-I did not reduce ADAM17 cell surface levels by the same magnitude (Figure 4B) (Lora et al., 2021), and LC viability was not affected (Figure 4C). The IFN-κ-driven inhibition of ADAM17 activity was abrogated by the JAK1/3 inhibitor tofacitinib without altering ADAM17 protein levels (Figure 4D-E) or cell viability (Figure 4F), supporting an IFNAR-mediated inhibition of LC ADAM17 sheddase activity rather than loss of ADAM17 cell surface expression. Application of IFN-κ to the skin of wild-type mice reduced LC ADAM17 activity by 57% (Figure 4G) while cell surface ADAM17 was reduced by 9% (Figure 4H). Together, these data indicate that IFN-I is sufficient to downregulate LC murine ADAM17 function and does so primarily at the level of sheddase activity ex vivo and in vivo.

**Figure 4.**
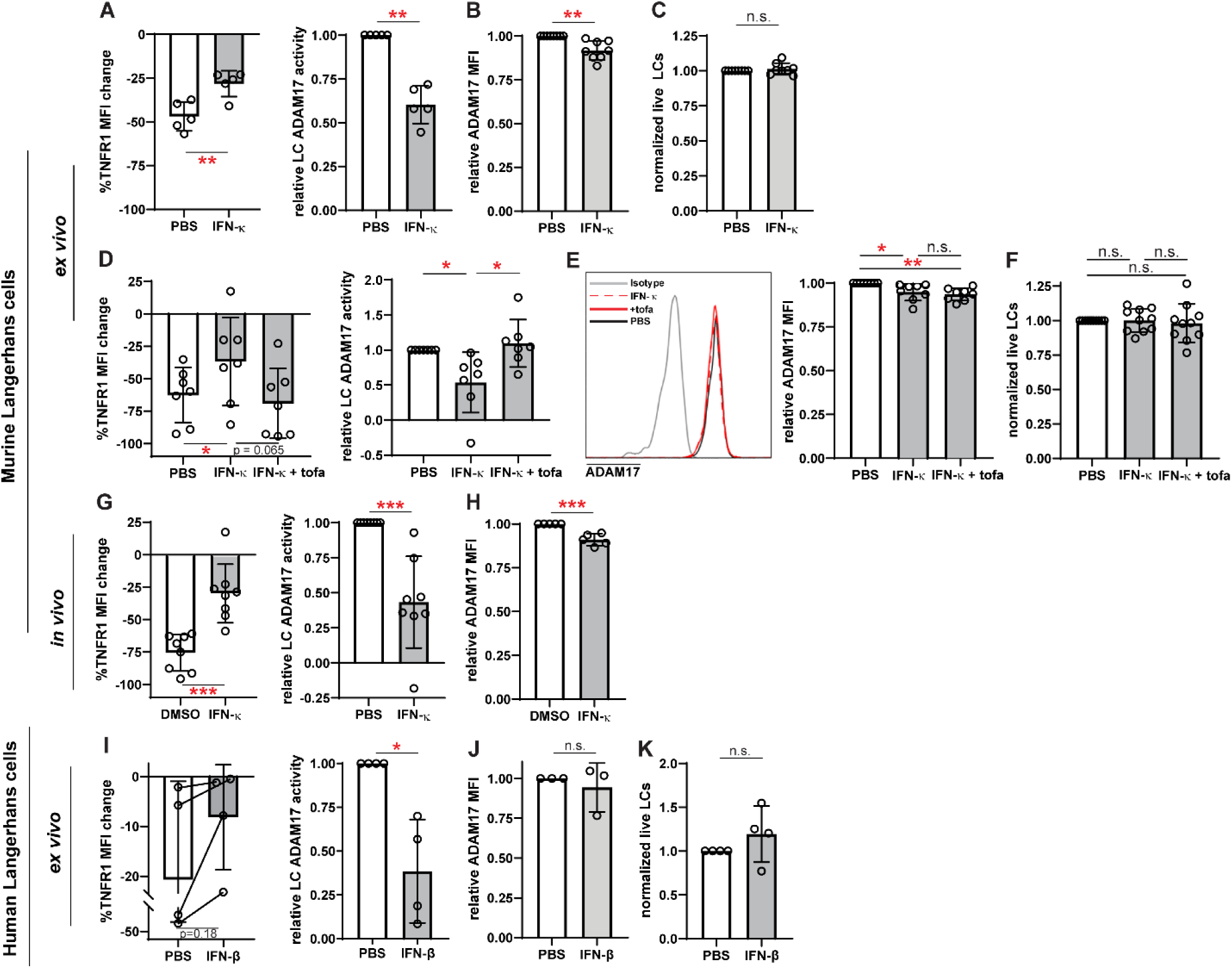
IFN-I inhibits LC ADAM17 sheddase activity. (A-K) Epidermal cell suspensions were treated ex vivo or mice were treated in vivo with IFN-I prior to assaying for UVR-induced LC ADAM17 sheddase activity, ADAM17 levels, and numbers. Cells from WT mice were treated with IFNκ or vehicle (A-C) with and without tofacitinib (D-F); IFN κ or vehicle was applied topically to ears of WT mice 16-20 hours prior to generation of epidermal cell suspensions (G-H); suction blister epidermal cell suspensions from healthy human donors were treated with IFNβ or vehicle (I-K). (A, D, G, I) LC ADAM17 sheddase activity as indicated by the extent of UVR-induced cell surface TNFR1 loss. Percent change in cell surface TNFR1 mean fluorescence intensity (MFI) after UVR (left); relative LC ADAM17 activity calculated by normalizing TNFR1 loss to that of vehicle controls (right). In (I), lines connect samples from the same donor. (B, E, H, J) Cell surface ADAM17 levels as indicated by MFI relative to that of vehicle controls. (C, F, K) Relative numbers of LCs per sample after indicated treatments. (A-K) Each symbol represents cells from a single mouse or donor, bars represent average values, and error bars are SD. n=3-10 per condition over 3-5 independent experiments. *p<0.05, **p<0.01, ***p<0.001, n.s.=not significant by unpaired t-test.

We also asked how IFN-I affects human LCs. While LCs can be obtained from skin biopsies or discarded tissues associated with surgical procedures, these tissues often require lengthy enzymatic digestion to dissociate the cells (Shipman et al., 2018), which has the potential to disrupt cell function. We found that epidermal roofs yielded from suction blistering could be digested for a short amount of time to obtain an epidermal cell mixture containing LCs (Figure S3) (Strassner et al., 2017). Addition of IFN-I reduced LC ADAM17 activity by 62% without altering ADAM17 expression or LC viability (Figure 4I-K). These results together showed that IFN-I can inhibit LC ADAM17 activity in both murine and human LCs and supported the idea that high levels of IFN-I could potentially contribute to LC ADAM17 dysfunction in disease.

### IFNAR is important for LC ADAM17 dysfunction in multiple lupus models

We next asked whether IFN-I was an important mediator of LC ADAM17 dysfunction in lupus models. Consistent with previous findings (Shipman et al., 2018), MRL/lpr mice showed a 42% reduction in LC ADAM17 sheddase activity when compared to control MRL/+ mice (Figure 5A). LC ADAM17 protein levels, however, were less affected, showing a 21% reduction (Figure 5B). Intraperitoneal (IP) anti-IFNAR treatment at 500 μg/dose restored LC ADAM17 activity in MRL/lpr mice (Figure 5A) without increasing ADAM17 protein expression (Figure 5B). A lower 125 μg/dose of anti-IFNAR had similar effects in rescuing LC ADAM17 sheddase function (Figure 5C-D). These results together suggested that IFN-I is important for inhibiting LC ADAM17 sheddase activity in MRL/lpr mice.

**Figure 5.**
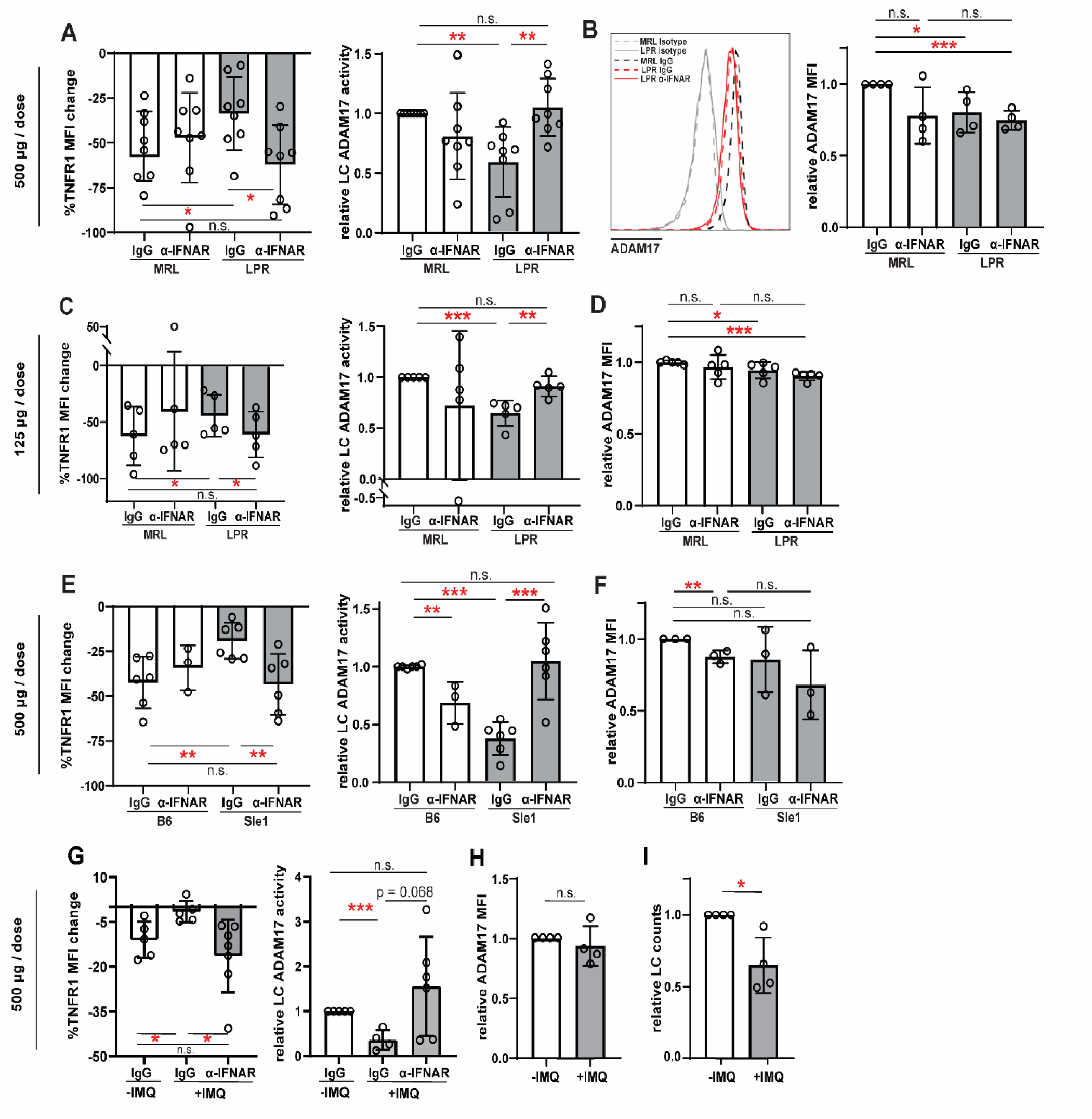
Anti-IFNAR restores LC ADAM17 activity in multiple lupus models. (A-I) MRL/lpr (A-D), B6.Sle1yaa (E-F), and IMQ (G-I) lupus model mice and their controls were treated twice with anti-IFNAR or isotype control at indicated doses over 6 days prior to collection of non-lesional epidermal cells. (A, C, E, G) UVR-induced LC ADAM17 sheddase activity as in Figure 4. (B, D, F, H) Relative cell surface ADAM17 levels. (I) Relative LC numbers per sample. (A-I) Each symbol represents a mouse, bars represent average values, and error bars are SD. n=3-8 per condition over 3-4 independent experiments. *p<0.05, **p<0.01, ***p<0.001, n.s.=not significant by paired (A, C, E (left)) or unpaired ((A, C, E (right)), B, D, F-I) t-test.

LCs from the B6.Sle1yaa model showed a 63% reduction in ADAM17 activity when compared to control mice, without a corresponding change in ADAM17 protein levels (Figure 5E-F). Similar to MRL/lpr mice, anti-IFNAR restored LC ADAM17 activity in B6.Sle1yaa mice without increasing ADAM17 protein expression (Figure 5E-F).

IMQ model mice showed reduced LC ADAM17 sheddase activity without altering cell surface ADAM17 protein levels (Figure 5G-H), similar to data from MRL/lpr and B6.Sle1yaa models. Interestingly, unlike the MRL/lpr and B6.Sle1yaa models where LC numbers were unchanged from controls (Shipman et al., 2018), skin LC numbers in the IMQ model were reduced by 35% (Figure 5I), echoing findings in human skin (Shipman et al., 2018). Similar to the MRL/lpr and B6.Sle1yaa models, anti-IFNAR restored LC ADAM17 activity in IMQ mice (Figure 5G). Together, the sufficiency of IFN-I in inhibiting LC ADAM17 activity and the restoration of LC ADAM17 activity by anti-IFNAR across 3 lupus models suggest that the IFN-I-rich environment drives LC ADAM17 dysfunction in non-lesional lupus skin.

### Anti-IFNAR reduces photosensitivity in an EGFR and LC ADAM17-dependent manner

We asked whether the anti-IFNAR-mediated restoration of LC ADAM17 function also reduced photosensitive skin responses. Anti-IFNAR treatment at 500 μg/dose reduced ear swelling in MRL/lpr mice (Figure 6A-B) but did not reduce epidermal permeability, a readout of skin barrier function (Figure 6C). Furthermore, in control MRL/+ mice, anti-IFNAR actually increased UVR-induced ear swelling and epidermal permeability (Figure 6B-C), suggesting that anti-IFNAR increased photosensitivity in non-lupus mice. These results echoed findings by other groups that IFNAR deficiency in non-lupus B6 mice increased photosensitivity (Sontheimer et al., 2017) and reduced skin wound re-epithelialization (Gregorio et al., 2010), suggesting that the 500 μg anti-IFNAR dose was disrupting intrinsic skin functions and potentially obscuring the effects of LC ADAM17 activity restoration. Consistent with this idea, anti-IFNAR at 125 μg/dose still worsened MRL/+ ear swelling but no longer increased epidermal permeability (Figure 6D-E), suggesting a decreased pathologic effect on homeostatic skin functions. This lower dose, which was sufficient to restore LC ADAM17 activity in MRL/lpr mice (Figure 5C), was sufficient to reduce both UVR-induced skin swelling and epidermal permeability (Figure 6D-E) and limit neutrophil and monocyte infiltration after UVR exposure (Figure 6F). At 125 μg/dose, then, anti-IFNAR rescued LC ADAM17 function and reduced UVR-induced skin swelling, epidermal permeability, and inflammatory infiltrates in MRL/lpr mice.

**Figure 6.**
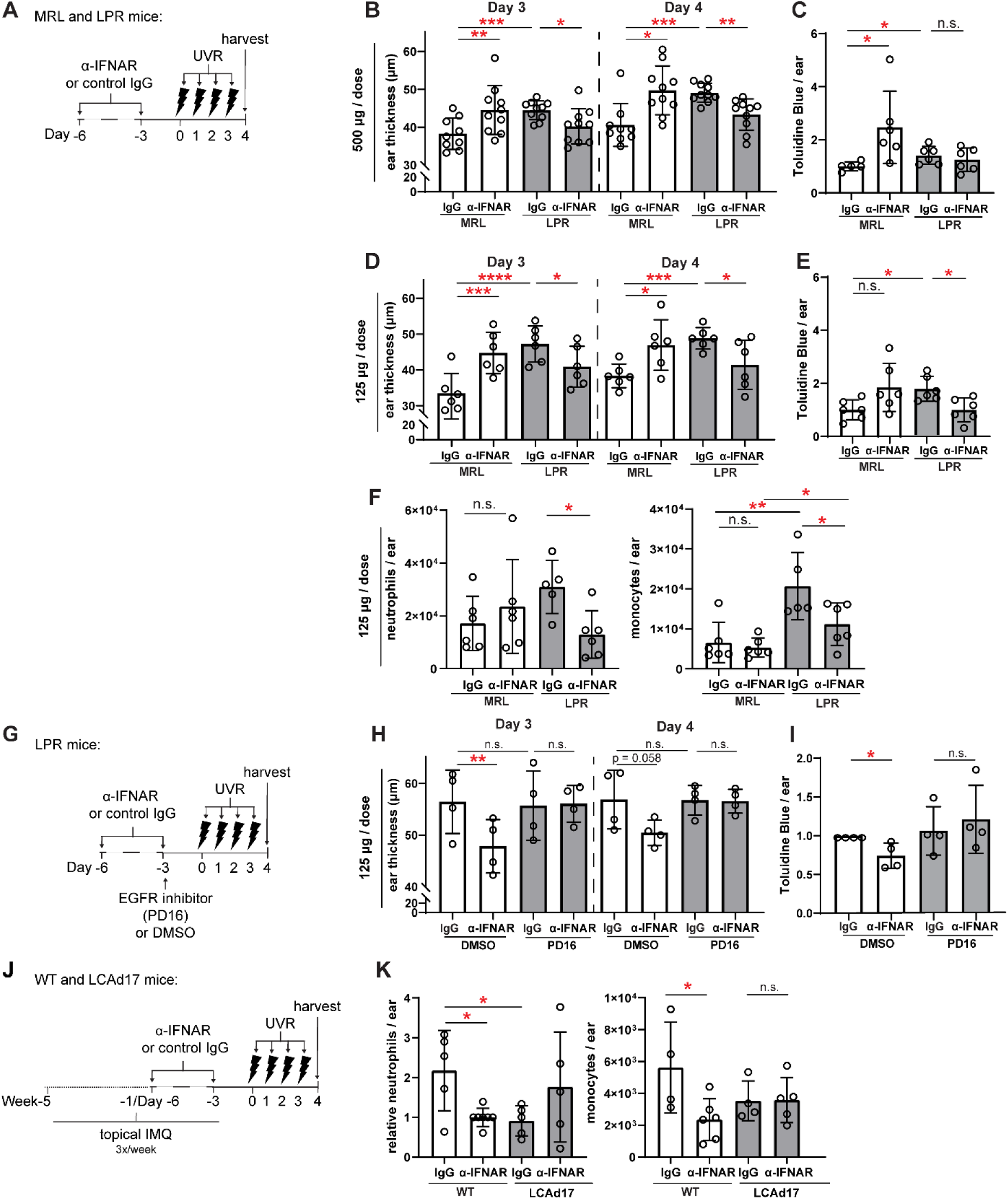
Anti-IFNAR reduces photosensitivity in EGFR- and LC ADAM17- dependent manners. (A-I) MRL/lpr and (J-K) LCAd17 mice and controls were treated according to schematics in (A, G, J) and non-lesional ears were harvested. (B, D, H) Ear thickness. (C, E, I) Epidermal permeability as indicated by toluidine blue retention. (F, K) Neutrophils and monocytes per ear. (B-F, H-I, K). n=4-10 per condition over 4-6 independent experiments. *p<0.05, **p<0.01, ***p<0.001, n.s.=not significant by paired (B, D, H) and unpaired (C, E, F, I, K) t-test.

We sought to understand the extent to which the anti-IFNAR-driven reduction in photosensitivity reflected the rescue of LC ADAM17 activity. As LC ADAM17 limits photosensitivity by stimulating EGFR signaling (Shipman et al., 2018), we blocked EGFR activity upon anti-IFNAR treatment of MRL/lpr mice (Figure 6G). EGFR blockade with the small molecule inhibitor PD16 abrogated the anti-IFNAR-mediated reduction in ear swelling and epidermal permeability (Figure 6H-I), suggesting that anti-IFNAR requires EGFR signaling to ameliorate photosensitivity in MRL/lpr mice. This is consistent with the idea that anti-IFNAR reduces photosensitivity by restoring LC ADAM17 function.

In the IMQ model, anti-IFNAR also reduced inflammatory cell infiltrate in UVR-treated skin (Figure 6J-K). In IMQ-treated LCAd17 mice lacking ADAM17 in LCs (Shipman et al., 2018), however, anti-IFNAR failed to reduce neutrophil and monocyte accumulation (Figure 6K), suggesting that the ameliorative effects of anti-IFNAR on UVR-induced inflammation are dependent on LC ADAM17. That anti-IFNAR reduced photosensitive responses across two lupus models depended on EGFR signaling and LC ADAM17 provides strong support for the idea that IFN-I contributes to photosensitivity at least in part by promoting LC ADAM17 dysfunction. These results also suggest that the beneficial effects of anifrolumab on SLE skin may be attributable to the improvement of LC ADAM17 activity.

### IFN-I inhibits UVR-induced LC ROS expression

UVR has been shown to trigger ADAM17 activity in fibroblasts via ROS generation (Singh et al., 2009), and we asked if IFN-I modulation of ROS was associated with changes in LC ADAM17 activity. We observed the expected UVR-induced ROS generation in LCs from WT skin (Figure 7A). Skin from IMQ-treated mice showed a reduction in UVR-induced LC ROS generation when compared to control skin, and anti-IFNAR treatment of IMQ mice restored UVR-driven ROS generation (Figure 7B). These results suggest that IFN-I could potentially limit UVR-induced LC ROS to inhibit LC ADAM17 function.

**Figure 7.**
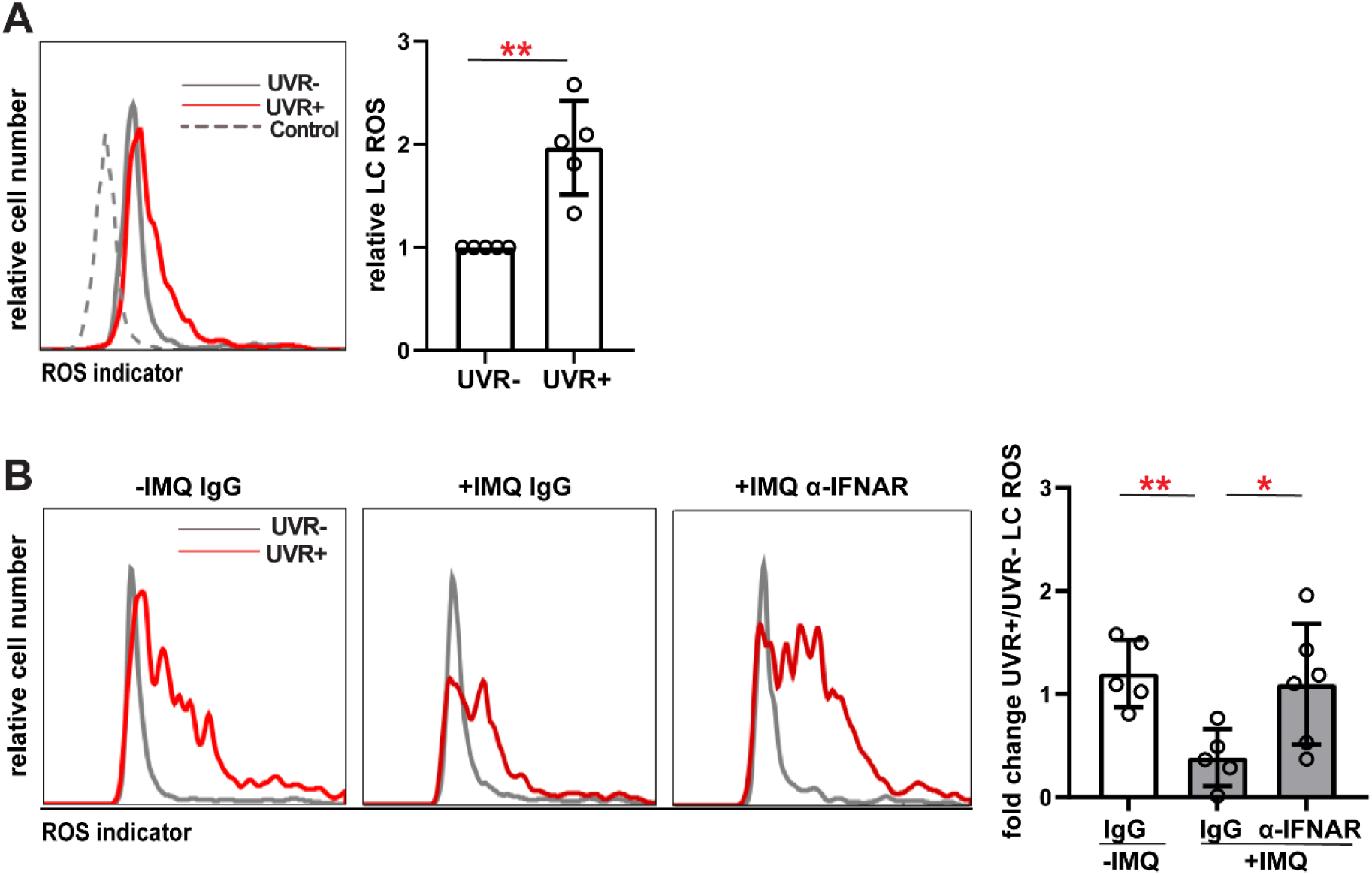
LC ROS is reduced in lupus model mice and restored by anti-IFNAR. (A-B) Epidermal cell suspensions from indicated mice were exposed to UVR or not and LC ROS content assayed by the ROS indicator CM-H2DCFDA. (A) WT mice; representative histograms (left) and LC ROS content relative to non-exposed condition (right). Dashed “control” line indicates signal without CM-H2DCFDA. (B) Control (-IMQ) and IMQ-treated mice (+IMQ) were treated as in Figure 5G with anti-IFNAR or isotype IgG prior to harvest of non-lesional skin; histograms (left) and fold change in LC ROS with UVR exposure (right). Each symbol represents one mouse, bars represent average values, and error bars are SD. n=5-6 per condition over 4-5 experiments. *p<0.05, **p<0.01, n.s.=not significant by unpaired t-test.

Together, our results suggested a model whereby the IFN-I-rich tissue microenvironment in lupus skin causes dysfunction of UVR-induced LC ADAM17, leading to photosensitivity. Anti-IFNAR restores this axis, suggesting that the therapeutic effects of anifrolumab on SLE skin could, in part, be via restoration of LC function.

## DISCUSSION

Here, we establish that multiple photosensitive lupus models resemble human lupus in showing upregulated IFN-I modules or IFN-regulated genes in non-lesional skin and that IFN-I in these lupus models causes LC ADAM17 dysfunction which contributes to photosensitive skin responses. These results suggest that the IFN-rich environment in lupus skin promotes photosensitivity by causing Langerhans cell ADAM17 dysfunction and that the beneficial effects of anifrolumab on skin disease in SLE patients, in part, reflects restoration of LC function. Our results also suggest that photosensitivity in the MRL/lpr, B6.Sle1yaa, and IMQ lupus models shares pathogenic mechanisms with human lupus, underscoring the utility of these models for the study of lupus skin pathophysiology.

How the IFN-I-rich environment in lupus skin contributes to photosensitive responses has not been well understood. Kahlenberg and colleagues have shown that IFN-I can potentiate IL-6 expression and apoptosis of human keratinocytes (Sarkar et al., 2018; Stannard et al., 2017). Keratinocytes are likely a major local source of IFN-I to modulate LC function in non-lesional skin, as LCs are positioned in the suprabasal layer of the epidermis near basal and spinous keratinocytes that demonstrate high IFN signaling (Billi et al., 2022). PDCs may also regulate LCs, as pDCs contribute to the IFN-rich environment in lupus skin (Furie et al., 2019; Karnell et al., 2021). Our results showing that anti-IFNAR restores LC ADAM17 function and reduces UVR-induced inflammation in an EGFR and LC ADAM17-dependent manner suggest that IFN-I contributes to photosensitivity by causing LC ADAM17 dysfunction. The accumulating evidence points to a pathogenic feed-forward loop between LCs and keratinocytes whereby keratinocyte IFN contributes to LC ADAM17 dysfunction and LC ADAM17 dysfunction contributes to keratinocyte dysfunction and ultimate loss. It should be noted, however, that the importance of IFN-I-mediated LC ADAM17 dysfunction does not rule out IFN-I-mediated keratinocyte-intrinsic mechanisms but points to multiple players in the ecosystem that maintain skin health.

Our findings showing that anti-IFNAR treatment could increase UVR-induced skin inflammation in control MRL/+ mice underscores the idea that excessive IFN-I contributes to skin pathogenesis in lupus. IFNAR is important for re-epithelialization during wound healing (Gregorio et al., 2010) and our data support the idea that our treatments should not compromise protective levels of IFNs. This notion is supported by the clinical findings that patients with high IFN-I scores responded to anifrolumab (Morand et al., 2020) and, when comparable, with a greater effect size than patients with lower IFN-I scores (Furie et al., 2017). Understanding the regulation and sources of the “excess” IFN-I in lupus will help to further develop therapies that ameliorate disease while preserving skin-intrinsic functions.

Our results extend our understanding of ADAM17 regulation as a contributor to disease. Increased ADAM17 activity can promote inflammation, such as by release of TNF or EGFR ligands within inflamed joints or lupus kidneys (Issuree et al., 2013; Qing et al., 2018) but loss of ADAM17 activity compromises barrier surfaces (Blaydon et al., 2011; Franzke et al., 2012).

Here we identify IFN-I as a negative regulator of ADAM17 in LCs and our data suggest a scenario whereby IFN-I reduces LC ROS generation to inhibit ADAM17 function. Further investigation will better delineate the locus and importance of LC ROS generation as well as contributions of other molecules such as the inactive Rhomboids (iRhoms) that regulate ADAM17 activity (Geesala et al., 2019).

High IFN-I states also include viral infections, genetic interferonopathies, and autoimmune diseases rheumatoid arthritis and dermatomyositis (Muskardin and Niewold, 2018), in which photosensitivity is found in the former and is a disease hallmark in the latter (Werth et al., 2004; Wysenbeek et al., 1989). Autoimmune disease-like manifestations that include photosensitivity are among the immune-related adverse events in cancer patients treated with checkpoint inhibitors (Brahmer et al., 2018), a situation where distinct IFN-I-mediated processes may have specific roles in promoting anti-tumor immunity and treatment resistance (Benci et al., 2019). Our findings will benefit patients with lupus and a wide spectrum of diseases.

## MATERIALS AND METHODS

### Study design

The purpose of the study was to understand how LC function is regulated by the environment in non-lesional skin to contribute to photosensitivity. The subjects are both humans and mice. We used gene expression analyses to understand the gene expression profiles in human lupus and murine lupus models. We used ex-vivo flow cytometry-based assays, assessment of epidermal permeability, and skin swelling assays to understand the effects of IFN on LC function and implications for photosensitivity. Sample sizes and replicate numbers are provided in each figure legend. Sample sizes were determined based on previous similar types of experiments. Mice were randomly assigned to treatment groups. No criteria were set for excluding certain data points and no data points were excluded. No specific confounder variables such as order of treatments or cage location were controlled for except that our experiments used multiple independent cohorts of mice over replicate experiments. Assessments and analyses were not blinded. ARRIVE reporting guidelines (Percie du Sert et al., 2020) have been used.

### Mice

8-16 week old mice were used unless otherwise specified. Mice were sex and age-matched. For C57Bl/6 mice, LCAd17 (Langerin-cre; Adam17 f/f) mice, and wildtype littermate controls, males and females were used. Unless otherwise indicated, for MRL/+ and MRL/lpr mice, females were used at 8-12 weeks, while males were used at 12-16 weeks as females present with a disease phenotype earlier than males. For B6.Sle1yaa mice and their B6 controls, only males were used because the model depends on the autoimmune accelerator locus located on the Y chromosome. B6 mice were either bred in-house or purchased from Jackson Laboratory (Bar Harbor, ME). MRL/+, MRL/lpr mice, B6.Sle1yaa mice, and their B6 controls were either bred in-house or purchased from Jackson Laboratory. LCAd17 mice (Shipman et al., 2018) were bred in-house. Mice were kept in a specific pathogen-free barrier facility and all animal procedures including protocols to reduce pain, suffering, and distress were performed in accordance with the regulations of the Institutional Animal Care and Use Committee at Weill Cornell Medicine (New York, NY).

### Mouse treatments

For IFN-κ painting, 37.5 ng IFN-κ (R&D Systems, Minneapolis, MN) was dissolved in DMSO (Sigma Aldrich, St. Louis, MO) and was applied topically onto each ear (18.75 ng on dorsal and ventral sides, each). For anti-IFNAR1 treatments, mice were injected IP with 500 µg/dose or 125 µg/dose of anti-IFNAR1 (MAR1-5A3) or isotype control (MOPC-21) (Bio X Cell, West Lebanon, NH). For UVR treatments, mice were exposed to 250 J/m2 (25 mJ/cm2)daily for four consecutive days using a bank of four FS40T12 sunlamps that emitted a combination of UVA and UVB radiation, similar to described (Shipman et al., 2018). Measurement using UVA and UVB meters revealed that these lamps emit 2.1mW/cm2 UVA and 0.8mW/cm2 UVB. To measure ear swelling after UVR exposure, a caliper (Mitutoyo, Aurora, IL) was used to take three measurements throughout each ear, and the reported value is the average of both ears. Each measurement was taken 22-24 hours after the previous measurement. For the IMQ-induced lupus mouse model, mice were painted on the dorsal and ventral sides of both ears with 5% imiquimod cream (42 mg/ear/mouse) 3x/week for 5-6 weeks (Yokogawa et al., 2014) prior to, and continued through experiments. Indicated mice received 2.95 mg of the irreversible EGFR inhibitor PD168393 (0.74 mg on dorsal and ventral sides, each) (Selleck Chem, Pittsburgh, PA).

### Human subjects

For microarray analysis, seven discoid CLE patients were examined. Both lesional and non-lesional skin were collected and the gene expression data from lesional tissue of these patients have been published, and patient characteristics are described (Jabbari et al., 2014). Healthy controls include the three that were previously described (Jabbari et al., 2014). All human tissue collection and research use adhered to protocols approved by the Institutional Review and Privacy Boards at the Rockefeller University (IRB# AJA-00740) and New York University (IRB# 10-02117), where participants signed written informed consents.

For suction blistering, 1 male and 3 female healthy participants between the ages of 23 and 55 participated. All human tissue collection and research use adhered to protocols approved by the Institutional Review and Privacy Board at the Hospital for Special Surgery (IRB# 2019- 1998), where participants signed written informed consents.

### Human skin cell collection and preparation

Suction blistering was performed as previously described (27). Eight 5 mm suction blisters were generated with a negative pressure instrument (NP-4 model, Electronic Diversities, Finksburg, MD) on the arm of healthy donors over 30-60 minutes. The blister fluid was then collected by aspiration, and epidermal roofs were collected. After collection, epidermal roofs from suction blisters were digested with dispase (2.42 U/mL; Thermo Fisher Scientific, Waltham, MA), collagenase type II (616 U/mL; Worthington Biochemical Corporation, Lakewood, NJ), and DNAseI (80 μg/mL; Sigma Aldrich, St. Louis, MO) for 40 minutes to generate single cell suspensions that were used for the ADAM17 sheddase assay.

### Skin cell collection and flow cytometry staining and quantification

For murine epidermal single cell suspensions, skin on the trunk was excised and subcutaneous fat scraped off. Skin was incubated in dispase (2.42 U/mL) at 37°C. The epidermis was then gently peeled, finely minced, digested in type II collagenase (616 U/mL) and DNAse I (80 μg/mL).

For collection of ear skin, ears were cut along the cartilage ridge, and the dorsal and ventral sides were manually peeled. Ear skin was then finely minced and digested with a buffer containing dispase, type II collagenase, and DNAse I as described above. For murine LC TNFR1 staining for the ADAM17 sheddase activity assay (see below), epidermal cells were preincubated with Fc block and then stained so that we could gate on CD45+, CD3-, CD11c+, I-Ab+ for B6 mice and I-Ak+ for MRL mice, and EpCAM+ LCs and assess for TNFR1. LCs were sometimes stained with rabbit-anti-ADAM17 (Lora et al., 2021) followed by donkey anti-rabbit IgG (Jackson Immunoresearch, West Grove, PA). Monocytes and T cells were identified by gating for CD45+, CD11b+, Ly6C+, and CD45+, CD11c-, I-Ak/I-Ab-, and CD3+ cells, respectively.

4’,6-diamidino-2-phenylindole dihydrochloride (DAPI) was used to exclude dead cells and debris (Sigma Aldrich, St. Louis, MO). All antibodies are from Biolegend, San Diego, CA unless otherwise indicated.

For human LC TNFR1 staining for the ADAM17 sheddase activity assay, single cell suspensions of the epidermal roofs of suction blisters and blister fluid were preincubated with Fc block and subsequently stained for CD45+, CD11c+, CD14-, CD3- CD19-, CD56-, CD66b-, EpCAM+, and HLA-DR+ to identify LCs. DAPI was used to exclude dead cells and debris, and TNFR1 was also stained to assess ADAM17 activity. All antibodies are from Biolegend, San Diego, CA unless otherwise indicated. Cells were analyzed using a FACS Canto or FACS Symphony A3 (BD Biosciences, San Jose, CA) and data were analyzed with FlowJo V10 Software (TreeStar, Ashland, OR).

For mean fluorescent intensity (MFI) calculations of TNFR1, each sample was assayed in triplicate, and the mean of the triplicates was recorded unless otherwise specified. The MFI of TNFR1 was divided by the MFI for the isotype control to generate a relative MFI. The percent TNFR1 MFI change was calculated by setting the relative MFI of the control sample to 1 and normalizing the value from the experimental samples to that control. In experiments with more than one control sample, the control values were averaged, and the experimental samples were each normalized to the averaged value.

For antibodies used, please see Table S9.

### Ex vivo ADAM17 sheddase activity assay

For murine skin, cell suspensions of epidermis were suspended at 200,000-500,000 cells/mL medium. For MRL/LPR and B6.Sle1yaa skin, medium was RPMI with 2% FBS, HEPES buffer, L-glutamine, penicillin-streptomycin (all Thermo Fisher Scientific, Waltham, MA). For IMQ mice skin, medium was phenol red-free RPMI (Corning, Edison, New Jersey), no serum, with HEPES buffer, L-glutamine, penicillin-streptomycin. Cells were exposed to 500 (LPR and B6.Sle1yaa model mice) or 250 (IMQ model mice) J/m2 UVR, similar to previous (Shipman et al., 2018), incubated for 45 minutes, and then stained for cell surface markers described above. For IFN-κ treatment, 3.125 ng/mL IFN-κ (R&D Systems, Minneapolis, MN) was added to the epidermal cell suspension that were incubated for 16-20 hours prior to UVR exposure. Indicated cells were treated with 1nM tofacitinib (Selleck Chem, Pittsburgh, PA) during IFN-κ incubation.

For human skin, cells were incubated for 1 hour with recombinant human IFN-β (35.7 ng/mL; activity=2.8×108 UI/mg) and then exposed to 500 J/m2 UVR as above. Cells were further incubated for 45 minutes and collected for preincubation with Fc block. Cells were then stained for cell surface markers described above.

### Epidermal permeability assay

Epidermal barrier function was assessed as described (Shipman et al., 2018). Ear skin was dehydrated and rehydrated in graded methanol and then incubated in 0.1% toluidine blue dye (Sigma Aldrich, St. Louis, MO). The dye was extracted with a buffer consisting of 95% methanol, 2.5% sulfuric acid, and 2.5% water and quantified with a colorimeter at 620 nm. Dye concentration was calculated using a standard curve.

### Ex vivo ROS assay

Epidermal cell suspensions in phenol red-free RPMI were exposed to 250 J/m2 UVR and incubated with the ROS indicator 5-(and-6)-chloromethyl-2’,7’-dichlorodihydrofluorescein diacetate, acetyl ester (CM-H2DCFDA) (Invitrogen, Waltham, MA) at 10uM for 45 minutes prior to cell surface marker staining for flow cytometry.

### RNA extraction

Female MRL/+ and MRL/lpr 12-14 weeks old and 12 week old B6 and B6.Sle1yaa male mice were used for RNA sequencing. RNA was extracted from ear skin using an RNEasy Kit (Qiagen, Germantown, MD) and quality confirmed on a BioAnalyzer 2100 (Agilent Technologies, Santa Clara, CA). For RT-qPCR of IMQ-treated mice, RNA was extracted with Trizol (Life Technologies) following the manufacturer’s protocol.

### RNA sequencing

Libraries were prepared with non-stranded and poly-A selection TruSeq RNA Sample Preparation kits. Sequencing libraries were sequenced by the Genomics Core Facility at Weill Cornell Medicine using a HiSeq 2500 (MRL/lpr, MRL/+ samples) and NovaSeq (B6, B6.Sle1yaa) at a depth of 15-30 million mappable paired-end reads per sample.

### Computational analyses of gene expression data

Microarray analysis was performed using healthy controls from an existing GEO dataset (GSE52471) obtained from GEO using GEOquery and combined with non-lesional skin sample data kindly provided by the authors of the original study (Jabbari et al., 2014). Microarray data were normalized with gcRMA normalization using the gcrma R package (Wu et al., 2021). Differential gene expression analysis was performed with limma using the empirical bayes method (Ritchie et al., 2015) and controlling for microarray hybridization kit and sex. Batch correction was performed using ComBat (Leek et al., 2021). Visualizations were generated using ComBat corrected data with plotly and ggplot2 in R.

For RNA sequencing analysis, read quality filtering and adapter trimming were performed using *fastp* (Chen et al., 2018). Filtered reads were mapped to the mouse genome (mm10), and exonic reads were counted against GENCODE release 27 annotation with the STAR aligner (Dobin et al., 2013) using a customized pipeline available athttps://gitlab.com/hssgenomics/pipelines. Differential gene expression analysis was performed with edgeR 3.30.2 using quasi-likelihood framework. Genes with low expression levels (<3 counts per million in at least one group) were filtered from all downstream analyses. P-values were adjusted to correct for multiple testing using the Benhamini-Hochberg FDR procedure. Genes with adjusted p-values < 0.01 were considered differentially expressed. Downstream analyses were performed in R using a Shiny-driven visualization platform (RNAseq DRaMA) developed at the HSS David Z. Rosensweig Genomics Research Center.

Differentially regulated pathways were identified using R QuSAGE 2.22.0 package (Yaari et al., 2013) with MSigDB C2 set (curated gene sets). All gene sets with less than 10 genes were excluded from the analysis. Pathways with Benhamini-Hochberg adjusted p-values <0.005 (non-lesional skin versus healthy control) or <0.05 (MRL/lpr versus MRL/+) were used for further analyses.

Ranked GSEA analysis was performed to define gene sets from MSigDB C2 enriched in human non-lessional skin and MRL/lpr mice. Log-transformed fold changes in relevant comparisons (non-lesional skin versus healthy control and MRL/lpr versus MRL/+) were used as ranking variables. Pathways with Benhamini-Hochberg adjusted p-values <0.05 were selected and the pathways’ similarity network was built to visualize gene overlap between pathways using the R/Bioconductor package *hypeR* (1.7.0) (Federico and Monti, 2020) and compared to enriched pathways in human SLE patients and mouse models.

The R/Bioconductor package, GSVA (Hänzelmann et al., 2013)(v1.36.3), was utilized as a non-parametric, unsupervised method to estimate enrichment of pre-defined gene sets in gene expression data from human DLE skin and non-lesional skin from MRL/lpr and B6.Sle1yaa mice. The inputs for GSVA were a matrix of expression values for all samples and curated gene sets describing select immune/tissue cell types and inflammatory cytokines. Low-intensity probes were filtered out if the interquartile range (IQR) of their expression values across all samples was not greater than 0. Enrichment scores were calculated using a Kolgomorov Smirnoff (KS)-like random walk statistic and represented the greatest deviation from zero for a particular sample and gene set. Scores across all samples were normalized to values between -1 (indicating no enrichment) and +1 (indicating enrichment). Gene sets used as input for GSVA are listed in Tables S3 and S7. The human gene sets were previously published (Martínez et al., 2022). The mouse plasma cell, T cell, myeloid cell, neutrophil, pDC, dendritic cell, and keratinocyte gene sets were derived from Mouse CellScan, a tool for the identification of cellular origin of mouse gene expression datasets. The mouse IFN gene sets have been previously described (Kingsmore et al., 2021). The mouse cytokine signatures were generated by an iterative process involving derivation through literature mining and GO term definitions provided by the Mouse Genome Informatics (MGI) GO Browser (Bult et al., 2019).

### qPCR

Both “lesional” and “non-lesional” ears were collected from each IMQ-treated and control mouse. Ears were flash-frozen with liquid nitrogen and stored at -80°C. Reverse transcription was performed using random primers and gene expression assessed by qPCR. PowerSYBR Green PCR Master Mix (Applied Biosystems) was used and assays were run on a QuantStudio 6 Pro (Applied Biosystems). Gene expression was normalized to the control for each target gene. Primers used in the qPCR assay are listed in Table S10.

### Statistical analyses

We determined the normality of data distribution using the Shapiro-Wilk test. For normally distributed data, we used two-tailed unpaired and paired Student’s t-tests for comparisons between two conditions to evaluate p-values as indicated. For data that were not normally distributed, the Mann-Whitney test was used for unpaired and Wilcoxon matched-pairs signed rank test for paired comparisons. Significance was defined as p<0.05. GraphPad Prism software was used. For figures showing normalized values, the control sample was set to one, and the experimental samples were normalized relative to the control for that experiment. For experiments that contained more than one control sample, the mean was obtained for the control samples, and the individual control and experimental samples were calculated relative to this mean.

### Data availability

The murine RNAseq data have been deposited in GEO (accession ID : GSE222573). All other data supporting the findings of this study are available within the paper and its Supplementary Materials.

## Supporting information

Table S10. Primer sequences IMQ RT-qPCR

Table S9. Antibodies used

Table S8. Ranked GSEA enriched pathways in human non-lessional skin

Table S7. Murine Gene Sets used for GVSA

Table S6. DEGs in B6.Sle1yaa vs control B6 non-lesional skin

Table S5. Pathways in LPR vs control MRL non-lesional skin using QuSAGE

Table S4. DEGs in LPR vs control MRL non-lesional skin

Table S3. Human Gene Sets used for GVSA

Table S2. Pathways in DLE vs healthy control non-lesional skin using QuSAGE

Table S1.DEGs in DLE vs healthy control non-lesional skin microarray

## ACKNOWLEDGEMENTS

**Acknowledgements:** The authors thank the members of the Lu Lab for helpful discussions and critical reading of the manuscript.

## Funding

Alpha Omega Alpha Carolyn L. Kuckein Student Research Fellowship (TML) Erwin Schrodinger Fellowship J 4638-B FWF (VZ)

National Institutes of Health grant T32AR071302-01 to the Hospital for Special Surgery Research Institute Rheumatology Training Program (NS and WDS)

National Institutes of Health grant MSTP T32GM007739 to the Weill Cornell/Rockefeller/Sloan-Kettering Tri-Institutional MD-PhD Program (WDS)

Tow Foundation (YC and DJO)

National Institutes of Health grant K08 AR069111 to the University of Iowa Department of Dermatology (AJ)

Veterans Administration VA Merit I01 BX004907 (AJ),

Physician Scientist Career Development Award from the Dermatology Foundation (AJ) National Institutes of Health grant R01AR077194 (AJ)

National Institutes of Health grant R01 DK099087 (IR) National Institutes of Health grant R35GM134907 (CPB) National Institutes of Health grant R01AI079178 (TTL) National Institutes of Health grant R21 AR081493 (TTL) Department of Defense grant W81XWH-21-LRP-IPA (TTL) Lupus Research Alliance (TTL)

St. Giles Foundation (TTL)

Barbara Volcker Center for Women and Rheumatic Diseases (TTL) A Lasting Mark Foundation (TTL)

National Institutes of Health Office of the Director grant S10OD019986 to Hospital for Special Surgery.

## AUTHOR CONTRIBUTIONS

Conceptualization: TML, TTL

Performed experiments and acquired data: TML, VZ, ESS, MD, KRV, NS, AJ

Data analyses and interpretation: TML, VZ, ESS, MD, YC, ARD, DJO, IR, CPB, PEL, TTL Methodology and reagents: JL, YL, WDS, WGA, JHZ, MR, NA, CPB

Supervision: SFT, KBO, JGK, IR, CPB, PEL, TTL

Writing—original draft: TML, VZ, MD, YC, AD, IR, PEL, TTL Writing—review and editing: All authors

## Competing interests

Andrea R. Daamen and Peter E. Lipsky are employees of AMPEL BioSolutions, but have no financial conflicts of interest to report. Jonathan H. Zippin is a stockholder in YouV labs, Hoth Therapeutics, and FoxWayne Inc and is a paid consultant for Pfizer and Hoth Therapeutics, Inc. Carl Blobel holds a patent on a method of identifying agents for combination with inhibitors of iRhoms. Carl Blobel and the Hospital for Special Surgery have identified iRhom2 inhibitors and have co-founded the start-up company SciRhom in Munich to commercialize these inhibitors.

Theresa T. Lu is a paid consultant for Pfizer.

## SUPPLEMENTARY MATERIALS

**Figure S1.** Further analyses of gene expression in human DLE and multiple murine lupus models.

**Figure S2.** ADAM17 expression levels in LCAd17 mice, UVR-induced cell surface TNFR1 loss on LCs in mixed epidermal cell suspensions is dependent on LC ADAM17.

**Figure S3.** Langerhans cell yield and gating with suction blistering of human skin.

**Table S1.** Differentially expressed genes in DLE vs healthy control non-lesional skin microarray

**Table S2.** Differentially expressed pathways in DLE vs healthy control non-lesional skin using QuSAGE

**Table S3.** Human gene sets used for GVSA

**Table S4.** Differentially expressed genes in MRL/lpr vs control MRL/+ non-lesional skin RNAseq

**Table S5.** Differential expressed pathways in MRL/lpr vs control MRL/+ non-lesional skin using QuSAGE

**Table S6.** Differentially expressed genes in B6.Sle1yaa vs control B6 non-lesional skin RNAseq

**Table S7.** Murine gene sets used for GVSA

**Table S8.** Ranked GSEA enriched pathways in DLE vs normal control dataset

**Table S9.** Antibodies used

**Table S10.** Primer sequences for qPCR

## Notes

https://www.ncbi.nlm.nih.gov/geo/query/acc.cgi?acc=GSE222573

https://www.ncbi.nlm.nih.gov/geo/query/acc.cgi?acc=GSE52471

### Summary of Updates

Revised abstract, additional methods, competing interest statements.

