## Supplementary figures for "The interferon-rich skin environment regulates Langerhans cell ADAM17 to promote photosensitivity in lupus"

**Figure S1.** Further analyses of gene expression in human DLE and multiple murine lupus models

**Figure S2.** ADAM17 expression levels in LCAd17 mice, UVR-induced cell surface TNFR1 loss on LCs in mixed epidermal cell suspensions is dependent on LC ADAM17

**Figure S3.** Langerhans cell yield and gating with suction blistering of human skin

**Table S1.** Differentially expressed genes in DLE vs healthy control non-lesional skin microarray

**Table S2.** Differentially expressed pathways in DLE vs healthy control non-lesional skin using QuSAGE

**Table S3.** Human gene sets used for GVSA

**Table S4.** Differentially expressed genes in MRL/lpr vs control MRL/+ non-lesional skin RNAseq

**Table S5.** Differential expressed pathways in MRL/lpr vs control MRL/+ non-lesional skin using QuSAGE

**Table S6.** Differentially expressed genes in B6.Sle1yaa vs control B6 non-lesional skin RNAseq

**Table S7.** Murine gene sets used for GVSA

**Table S8.** Ranked GSEA enriched pathways in DLE vs normal control dataset

**Table S9.** Antibodies used

**Table S10.** Primer sequences for qPCR

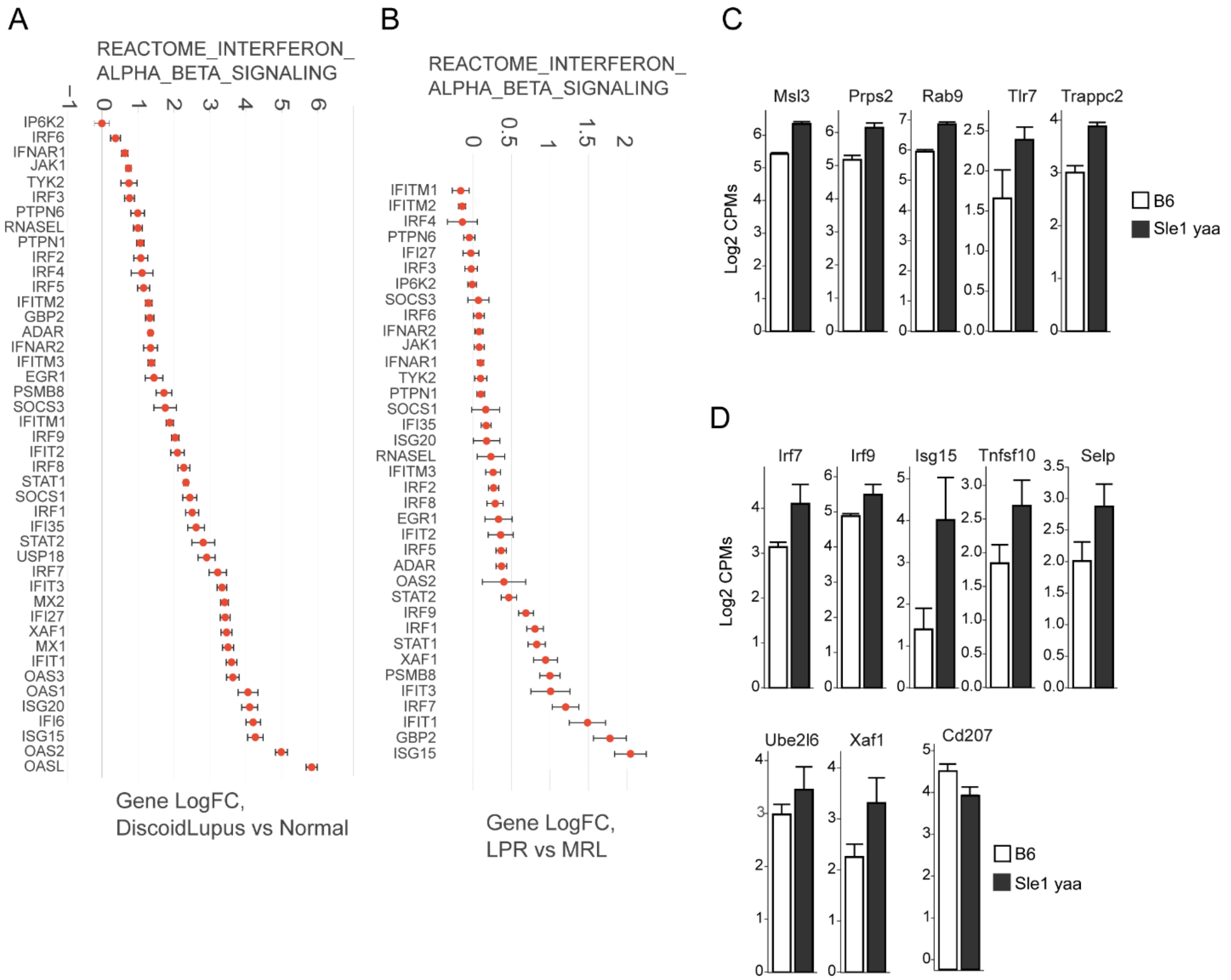


**Figure S1. Further analysis of gene expression in human CLE and multiple murine lupus models.** (A-B) Log-transformed expression fold change for genes in the IFN α/β pathway for human CLE and MRL/lpr model mice. (C) Genes from yaa locus are expressed at a higher levels in B6.Sle1yaa mice (D) IRF transcription factors and their targets are expressed at a higher levels and CD207 is expressed at lower level in B6.Sle1yaa mice compared to controls.

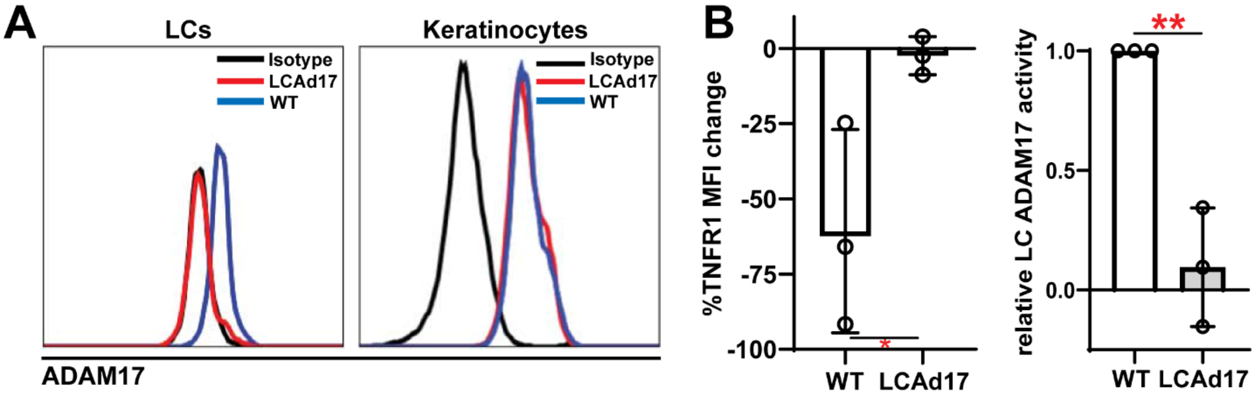


**Figure S2. ADAM17 expression levels in LCAd17 mice, UVR-induced cell surface TNFR1 loss on LCs in mixed epidermal cell suspensions is dependent on LC ADAM17.**
(A-B) Epidermal cell suspensions from WT or LCAd17 mice were examined. (A) ADAM17 cell surface protein expression on LCs and keratinocytes. Representative of 3 experiments. (B) UVR-induced cell surface TNFR1 loss on LCs is dependent on LC ADAM17. (Left) % reduction in cell surface TNFR1 mean fluorescence intensity (MFI). (Right) LC ADAM17 activity is plotted as a function of the relative reduction in TNFR1 loss, normalized to the change in WT mice. Error bars are SD. **p<.01 , n.s.=not significant by unpaired t-test.

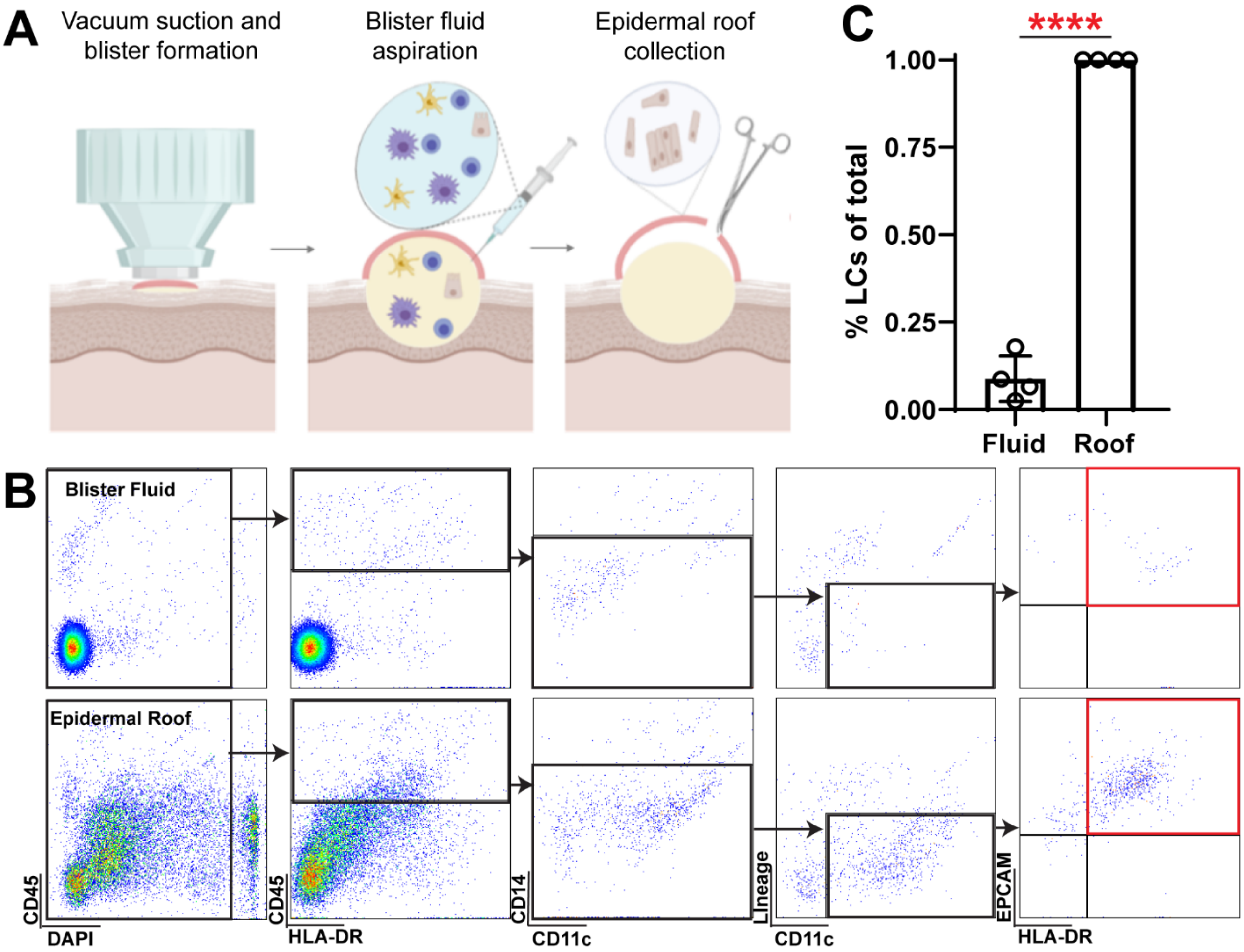


**Figure S3. Langerhans cell yield and gating with suction blistering of human skin.** (A) Schematic of suction blistering and sample collection. (B) Gating of Langerhans cells in blister fluid and in epidermal roof cell suspension. (C) Langerhans cell abundance in blister fluid and epidermal roof samples. Each symbol represents 1 healthy donor. ****p<.0001 unpaired t-test.
